# Theoretical understanding of evolutionary dynamics on inhomogeneous networks

**DOI:** 10.1101/2023.02.02.526861

**Authors:** Hamid Teimouri, Dorsa Sattari Khavas, Cade Spaulding, Christopher Li, Anatoly B. Kolomeisky

## Abstract

Evolution is the main feature of all biological systems that allows populations to change their characteristics over successive generations. A powerful approach to understand evolutionary dynamics is to investigate fixation probabilities and fixation times of novel mutations on networks that mimic biological populations. It is now well established that the structure of such networks can have dramatic effects on evolutionary dynamics. In particular, there are population structures that might amplify the fixation probabilities while simultaneously delaying the fixation events. However, the microscopic origins of such complex evolutionary dynamics remain not well understood. We present here a theoretical investigation of the microscopic mechanisms of mutation fixation processes on inhomogeneous networks. It views evolutionary dynamics as a set of stochastic transitions between discrete states specified by different numbers of mutated cells. By specifically considering star networks, we obtain a comprehensive description of evolutionary dynamics. Our approach allows us to employ physics-inspired free-energy landscape arguments to explain the observed trends in fixation times and fixation probabilities, providing a better microscopic understanding of evolutionary dynamics in complex systems.

## 1 Introduction

The most unique property of all biological systems is their ability to evolve over time by preferentially selecting randomly appearing features that benefit them most [8, 14]. While the main trends of evolution are now reasonably well understood, many aspects of evolutionary dynamics remain unclarified [8, 17]. In recent years, it was proposed to explore evolutionary dynamics on graphs as a way to mimic evolutionary processes for populations that possess complex structures, for example, as typically found in biological tissues [13, 29]. This approach has been widely utilized for investigating a variety of phenomena ranging from cancer initiation and evolution to social cooperativity and ecological dynamics, providing new insights into mechanisms of these processes [9, 11, 19, 20, 23–26, 32]. There have been multiple observations confirming that spatial structure of populations might have a strong effect on evolutionary dynamics [2, 4, 9, 12, 15, 18], but there is still no clear understanding of why it is happening.

It is widely accepted that populations evolve following a specific sequence of events [17]. After a random mutation appears in one of the individuals in the population, it might proliferate in the system via selection and random drift, eventually spreading to the whole population in a process known as fixation. But the fixation is not guaranteed, and the mutation might also disappear since the selection processes in the successive generations are random. Then the most crucial properties to characterize these processes are a fixation probability, which is defined as the probability for the given mutation to fully occupy the population, and a fixation time, which is defined as the mean time between the first appearance of the given mutation and its final fixation [1, 10, 22, 26, 27, 29, 31]. Because biological systems are typically very inhomogeneous, the fixation processes in these systems have been frequently investigated by exploring methods of evolutionary dynamics on graphs, which led to several remarkable observations [13, 26, 27, 29]. For example, while it was naively expected that homogeneous well-mixed populations exhibit the highest fixation probabilities, several network topologies have been identified expressing even higher fixation probabilities [13, 27, 29]. These systems have been labeled as amplifiers, and it has been suggested that they might accelerate the evolution [13]. However, all these networks amplify the selection of mutations at the cost of significantly slowing down the fixation dynamics, i.e., the fixation times in these systems are always larger than the fixation times for similar-size homogeneous well-mixed populations [28]. Although the fixation processes for inhomogeneous populations have been intensively studied in recent years [3, 5, 6, 16, 21, 27, 29], there is still no clear understanding on the microscopic origin of selection amplifications, the connections to the underlying network topology, and the correlations between the fixation probabilities and the fixation times.

In this paper, we present a theoretical investigation of evolutionary dynamics on inhomogeneous populations by applying a method of stochastic mapping [26]. In this approach, evolutionary changes in the system can be viewed as stochastic transitions between discrete states that are specified by different numbers of mutated individuals in the populations. To be more specific, we explicitly analyze evolutionary dynamics on star networks that are known to be selection amplifiers. Explicit expressions for fixation probabilities and fixation times are obtained using first-passage probabilities calculations and physically consistent approximations. Theoretical calculations are supported by extensive Monte Carlo computer simulations. It is argued that the overall evolutionary process in the system can be viewed as a motion in the effective free-energy landscape, allowing us to explain the microscopic origin of amplification and the observations of larger fixation probabilities together with slower fixation times.

## 2 Theoretical Method

### 2.1 Evolutionary Dynamics on Graphs

Let us investigate a specific biological population that can evolve following random mutations and sequential selection processes. To be specific, we consider an originally healthy tissue with *N* wild-type stem cells (i.e., those cells that can replicate). At some time (assumed to be *t* = 0), a mutation appears in one of the cells [7, 30]. The tissue cells can replicate, although the rates are different for normal (wild-type) and mutated cells. It is assumed that the division rate for normal cells is equal to *b*, while the mutated cells dive with a rate *r* × *b*. The parameter *r* is defined here as a fitness parameter that specifies how faster is the replication rate for the mutated cells in comparison with the wild-type cells. It plays a critical role in dynamic processes since it assists the evolution in choosing the specific mutations to take over the whole tissue [17, 26]. For *r* > 1, the mutations are viewed as advantageous, while for *r* < 1 the mutations are disadvantageous. In addition, *r* =1 specifies neutral mutations. For convenience, we assume here that the replication rate of normal cells is *b* = 1.

Another crucial factor that drives the evolution is a requirement to have the total number of cells *N* to be constant [26]. For biological tissues, it is a consequence of homeostasis when the most relevant physiological properties of organisms tend to be constant [14]. Although the specific mechanisms of how the number of cells in the tissues are kept constant at the microscopic level are not yet fully understood, the popular approach to mimic the processes that support the homeostasis is to utilize a so-called Moran procedure [17]. It is a two-step process. First, one of *N* cells is randomly chosen to replicate proportionally to its fitness. This temporarily increases the number of cells in the tissue to *N* + 1. Then one of *N* + 1 cells is chosen to be instantaneously removed to return to the original number of cells in the tissue.

To better understand the complex dynamic processes in biological systems, it has been proposed to investigate the evolutionary dynamics on networks [13, 29]. This is schematically illustrated in Fig. 1. The idea here is that networks efficiently reflect spatial inhomogeneity and variations in activity in the biological tissues. In this approach, each node corresponds to one cell, and connections between nodes specify the direction of selection processes after the replication. The advantage of analyzing the dynamics on graphs is that both homogeneous (Fig. 1a) and inhomogeneous networks (Fig. 1b) can be investigated in one framework, allowing for better understanding of the role of population structures in evolutionary dynamics.

**Figure 1.**
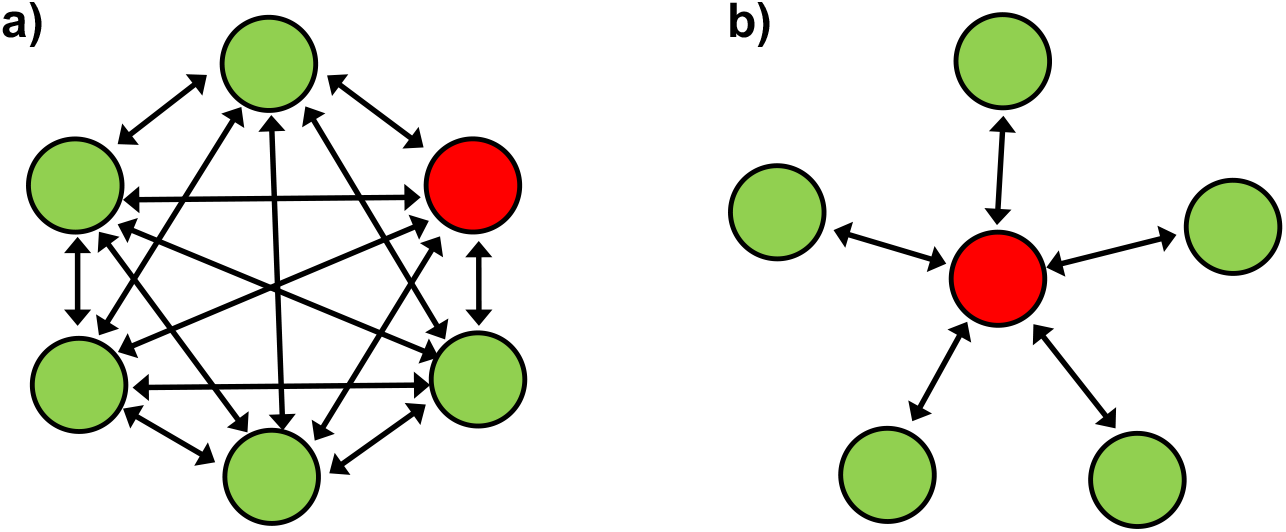
Schematic view of evolutionary dynamics on networks. Arrows indicate allowed changes after the replications. Green cells are normal and red cells are mutated. a) A homogeneous well-mixed model of size *N* where the replications at any node can lead to the removal of any other (*N* – 1) cells with equal probability 1/(*N* – 1). b) An inhomogeneous network where the replication at one special star node can affect any other (*N* – 1) cells with probability 1/(*N* – 1), while the replications at any of the (*N* – 1) branched cells can only change the star node with unit probability.

To explain evolutionary processes on graphs, let us first consider the homogeneous network presented in Fig. 1a. In this model, there are *N* identical cells, and the Moran procedure here is the following.

After the randomly selected cell replicates, temporary increasing the number of cells to *N* +1, with the probability 1/(*N* – 1) any other of (*N* – 1) cells is substituted by the newly created cell, bringing down the number of cells again to N. This is a well-mixed homogeneous system for which the fixation dynamics has been fully investigated [13, 17, 26]. More specifically, the fixation probability for this system is equal to

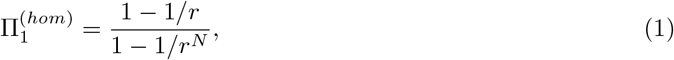

while the fixation time is given by [24],

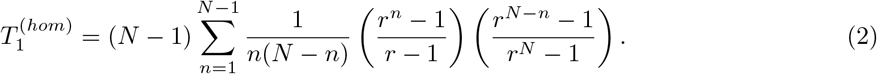

The sub-index “1” in these expressions corresponds to the fact that the evolutionary process starts with just one mutated cell, while the super-index *hom* reflects that the system is homogeneous. In the limit of *r* → 1 (neutral mutations), the fixation probabilities and fixations times simplify into

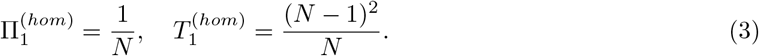

Another important limiting and more realistic case is when *N* → ∞ for *r* > 1. In this case, it can be shown that

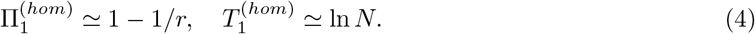

### 2.2 Evolutionary Dynamics on Star Networks

Now let us consider the evolutionary dynamics for inhomogeneous populations. More specifically, we concentrate on the star network as presented in Fig. 1b. This is the system where the fixation probability amplifications has been observed [13, 28]. In this model, there are two types of cells: one central node and (*N* – 1) branched nodes. After the replication takes place in the central cell, the selection can substitute any of the branched cells with the probability 1/(*N* – 1). However, if the replication occurs at the branched cells, then only the central cell will be substituted with the unit probability.

To investigate the fixation dynamics in the star network, we explore a method of stochastic mapping that has been already successfully utilized for understanding cancer initiation processes [23—26]. The main idea of this approach is to view the evolutionary processes as a set of stochastic transitions between different states. These states are specified by the number of mutated cells. For the star model, this approach is illustrated in Fig. 2. We define the state *n* as the one that has *n* mutated cells, but only in the branched nodes and not in the center, while the state *n*^(0)^ defines the situation with *n* mutated cells that includes the central node. Arrows in Fig. 2 identify possible transitions between the states. There are four types of transition rates. The rate 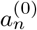 describes the transitions from the state *n*^(0)^ to the state *n* + 1, and the rate *a_n_* describes the transition from the state *n* to the state *n* + 1 (Fig. 2b). It corresponds to the increase in the number of mutated cells in the system. The decrease in the number of mutated cells are given by the rates 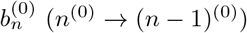 and the rates *b_n_* (*n* → (*N* – 1)^(0)^): see Fig. 2b. Importantly, one can see two chains of states between the state without mutations (*n* = 0) and the fully mutated state (*n* = *N*): see Fig. 2. But effectively only one of them leads to the fixation.

**Figure 2.**
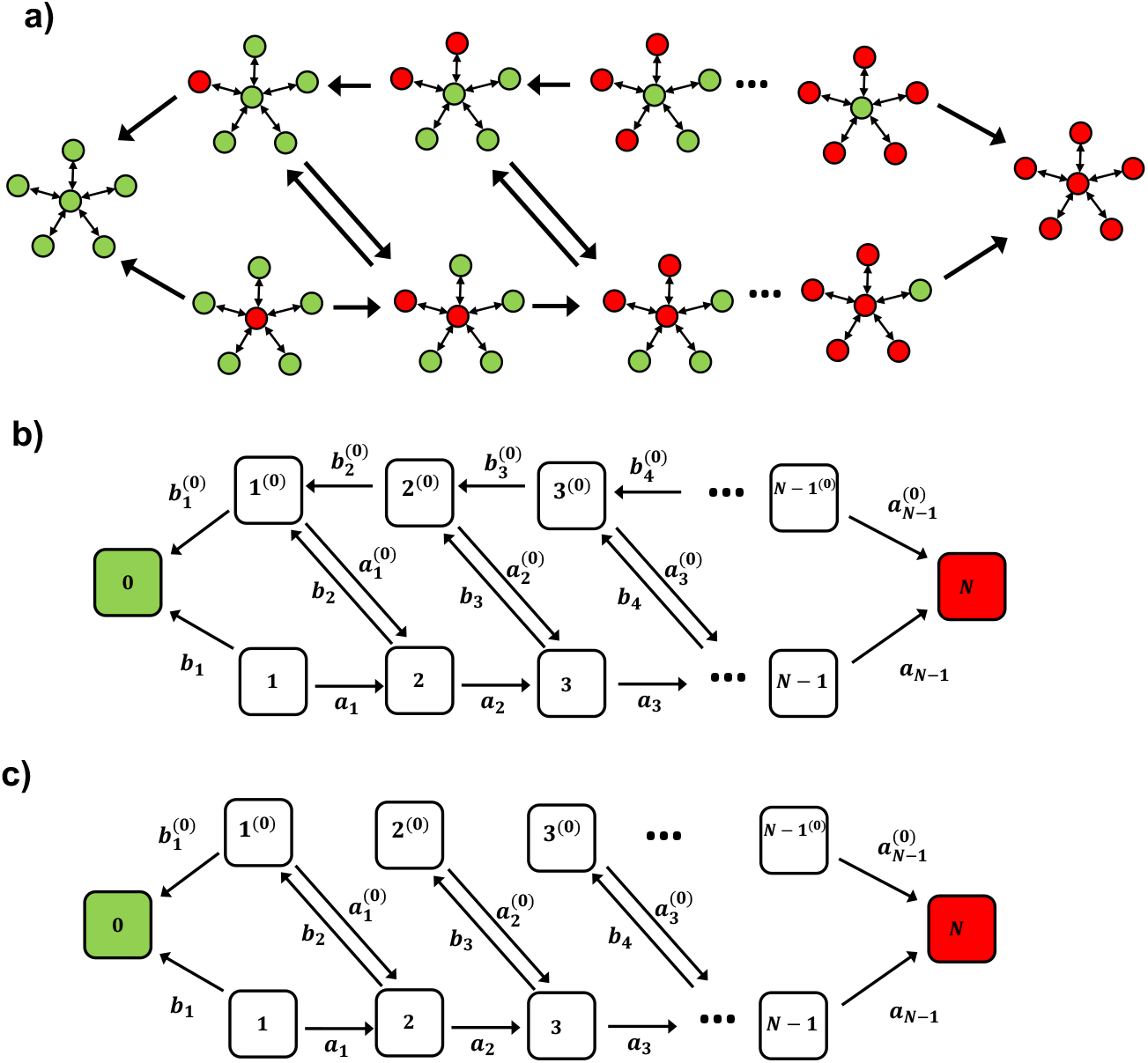
a) Evolutionary dynamics on the star network as a set of stochastic transitions. Green nodes correspond to normal cells while red nodes describe the mutated cells. Arrows correspond to allowed transitions. b) Corresponding discrete-state stochastic scheme of evolutionary dynamics for the star-model. State 0 describes the full elimination of all mutations, while state *N* corresponds to fixation c) An approximate model that neglects the reverse transition from the state *n*^(0)^ to the state (*N* – 1)^(0)^ for *n* > 1 - see text for more details.

As explained in the Supporting Information, the specific expressions for the transition rates are given by

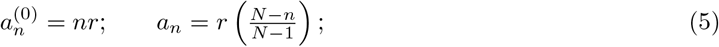

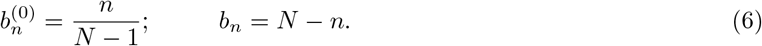

This allows us to fully evaluate the fixation dynamics on the star networks. For this purpose, we utilize a method of first-passage probabilities that has been successful in analyzing the mechanisms of cancer initiation [24–26]. We define functions 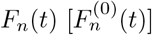 as the probability densities of reaching the fixation state *n* = *N* at time *t* if at *t* = 0 the system started in the state *n* [*n*^(0)^]. The time evolution of these probability density functions is governed by the following set of backward master equations,

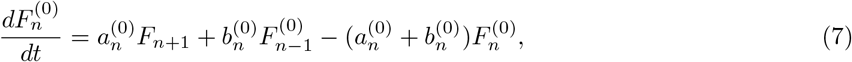

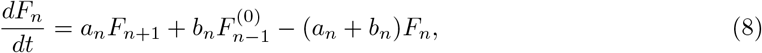

with initial condition *F_N_*(*t*) = *δ*(*t*), which means that if the system starts in the state *n* = *N* the fixation is immediately accomplished.

From the first-passage probabilities, the details of evolutionary dynamics on star networks can be fully identified. More specifically, one can calculate the fixation probabilities 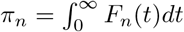 and the fixation times 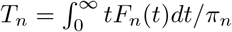. As shown in the Supporting Information, the fixation probabilities starting from the states *n* or *n*^(0)^ are given by,

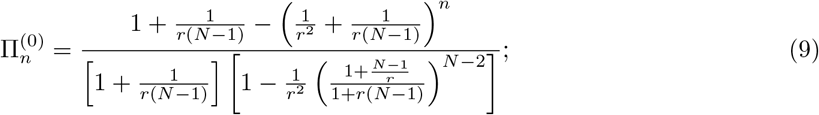

and

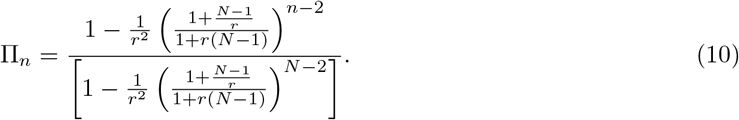

Interestingly, it can be shown that generally for all values of *n* we have 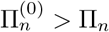. If starting from the single-mutation states (*n* = 1), the fixation probabilities are equal to

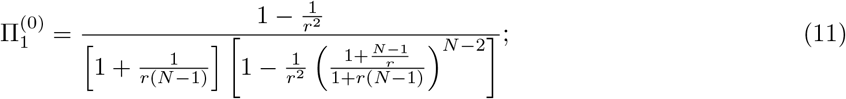

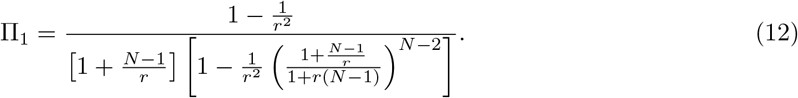

To better understand the microscopic picture of fixation processes, it is useful to consider limiting situations. If the replication rates of mutated cells are the same as for the normal cells (*r* = 1, neutral mutations), from Eqs. (11) we obtain

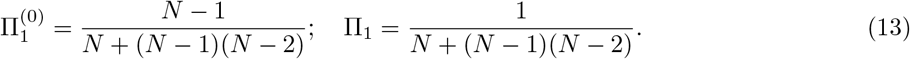

which for large number of cells (*N* ≫ 1) simplify into

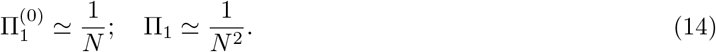

Another important limit is when *r* > 1 and *N* → ∞. In this case, it can be shown that

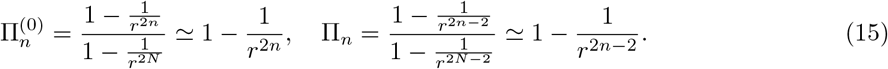

Starting from the single-mutation states (*n* = 1), these calculations yield

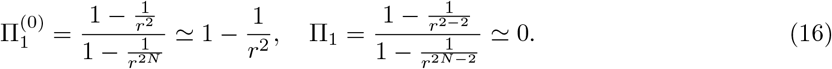

When the mutation appears first in one of the branched cells (starting in the state 1^(0)^) our limiting results fully agrees with previous calculations for the fixation probability in star networks [13]. But starting in the center of the network (the state 1) does not essentially lead to the fixation at all for large *N*. The results of our calculations are presented in Fig. 3. The most interesting observation here is that the fixation probability strongly depends on which initial cell is mutated. The mutation in one of the branched states leads to the amplification of fixation probabilities, while the mutation in the center node of the network significantly decreases the probability of fixation: see Fig. 3. Our theoretical approach allows us to clearly understand these observations. From the discrete-state stochastic scheme in Fig. 2b, one might conclude that the probability of eliminating the mutation from the state 1^(0)^ is given by

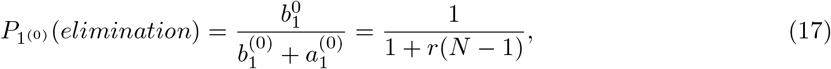

which in the limit *N* → ∞ approaches zero. The probability of eliminating the mutation from the state 1 is given by

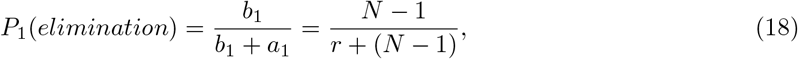

which in the limit *N* → ∞ approaches unity. Since the fixation is opposite to the mutation elimination [Π_*j*_ = 1 – *P_j_*(*elimination*)], one can see that it is more probable to remove the mutation from the central cell, while it is much less probable to eliminate the mutation from the branched cell. This is the origin of the fixation amplification phenomenon in the star networks.

**Figure 3.**
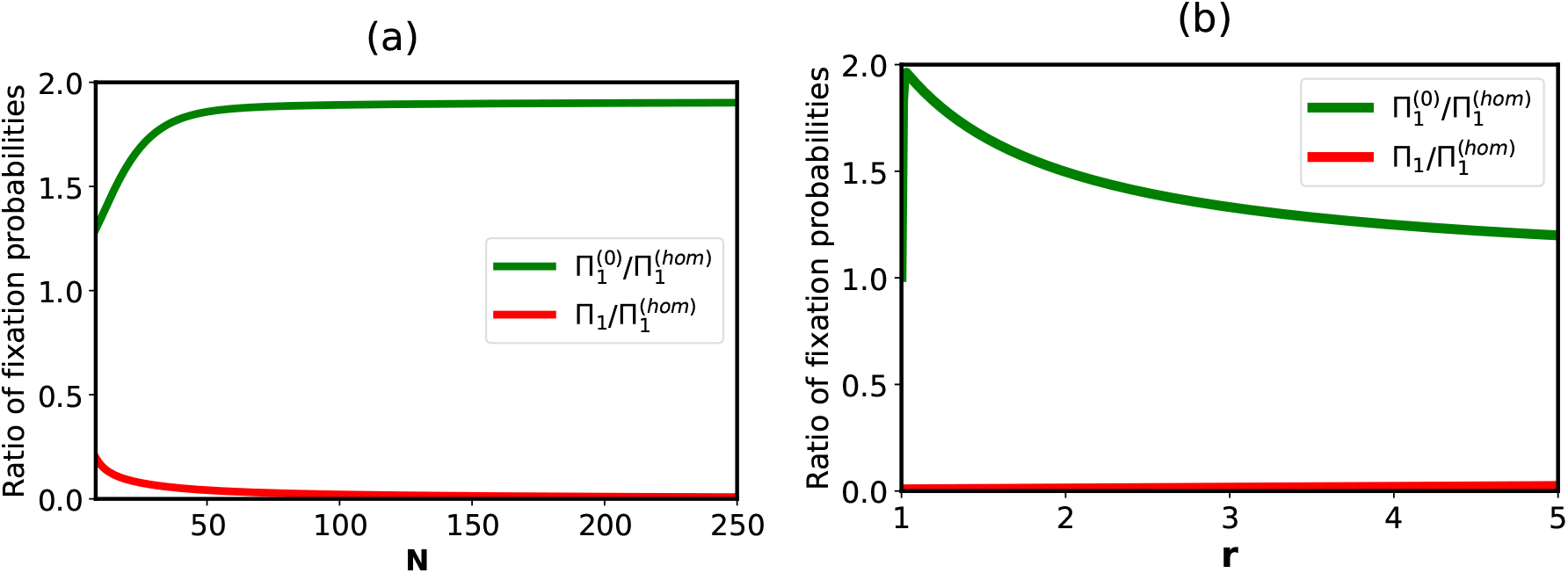
Ratio of fixation probabilities for the star network and the well-mixed homogeneous system: a) as a function of the system size for *r* =1.1, and b) as a function of the fitness parameter *r* for *N* = 250.

The results presented in Fig. 3b also suggest that the highest degree of amplification 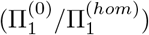 cannot be larger than two, and it can be achieved only for the fitness parameters that are only slightly larger than one. Interestingly, for neutral mutations (*r* = 1), our theoretical calculations predict that there will be no amplification at all 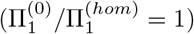. This is a consequence of the behavior of fixation probabilities at *r* =1 and large *N*, namely 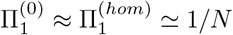. Thus, the fixation amplification works well only for slightly advantageous mutations.

Another interesting observation from our theoretical analysis is the dependence of the degree of amplification on the system size (Fig. 3a) and the fitness parameter *r* (Fig. 3b). Increasing the number of cells in the tissue makes the amplification stronger. This is because the probability of mutation elimination from the state 1^(0)^ behaves as 1/*N*. Surprisingly, making the mutation more advantageous (*larger r*) lowers the degree of amplification - see Fig. 3b. It can be shown that 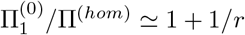 for large number of cells. This can be explained by arguing that there are more pathways to reach the fixation in the well-mixed homogeneous system, while it is only one pathway in the star-network model. Then, larger fitness parameters *r* increase the fixation probability more for the homogeneous system than for the inhomogeneous system.

While we were not able to obtain explicit expressions for the fixation times, they can be evaluated numerically by solving the corresponding backward master equations, as shown in the Supporting Information. In addition, we also run Monte Carlo computer simulations to evaluate the fixation dynamics in the star networks. The results of our numerical calculations and computer simulations are presented in Fig. 4 and compared with the homogeneous well-mixed model. Only the fixation times from the state 1^(0)^ are presented there because they are essentially the same as the fixation times starting from the state 1. One can see that the ratio of fixation times grows linearly with the size of the system (Fig. 4a), suggesting that for *N* ≫ 1 the fixation time on the star network scales as *T*_1(0)_ ~ *N* ln *N*. This can be explained in terms of our method of stochastic mapping. In the inhomogeneous star model there is one pathway that leads to the fixation - the one when the center cell is always mutated. However, in the well-mixed homogeneous system there are *N* such pathways that lead to the fixation since there are no topological constraints there. Fig. 4b shows that increasing the fitness advantage of the mutated cells accelerates the fixation dynamics in the star network, but the effect is rather modest. We also found that the slowest fixation dynamics is observed for very slightly advantageous mutations where the the fixation amplification is the strongest.

**Figure 4.**
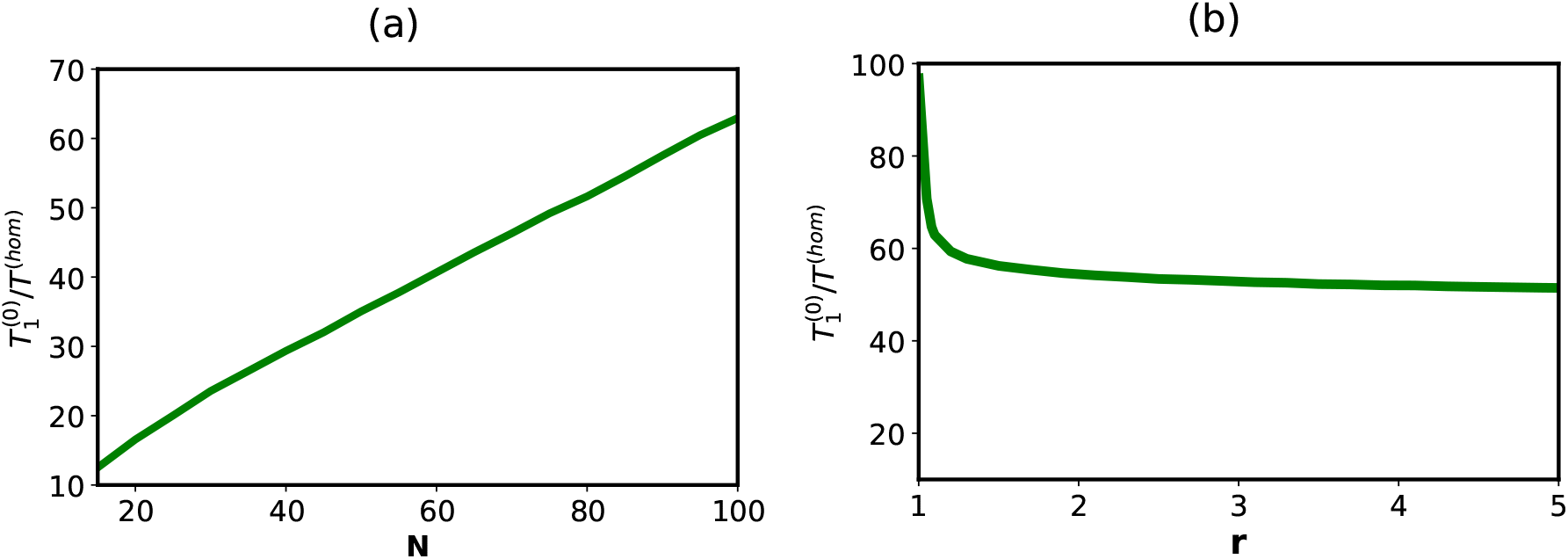
Ratio of fixation times for the star network and the well-mixed homogeneous system: a) as a function of the system size for *r* = 1.1, and b) as a function of the fitness parameter *r* for *N* = 100.

Our discrete-state stochastic description allows us to better understand the microscopic mechanisms of evolutionary processes on star networks. It shows that the amplification of the fixation probability is taking place not due to increased number of pathways to reach the fixation state but by lowering the probability of mutation elimination from the branched cells. However, it cannot lead to faster fixation dynamics because the system is frequently trapped in the states where the central node is not mutated (see Fig. 2a), slowing the overall fixation dynamics. In addition, the topology of the network dictates that there is only one pathway to reach the fixation state, while there are many more opportunities in the well-mixed homogeneous systems of the same size to reach the fixation state.

### 2.3 Approximate Model to Describe Evolutionary Dynamics on Star Networks

To better illustrate the idea of slowing the fixation dynamics in the star network due to trapping the system in the unproductive states, we propose considering an approximate model that captures main features of the evolutionary dynamics on star networks and allows us to obtain the explicit expressions for the fixation probabilities and fixation times. Our idea here is based on the observation that the probability to move from the state *n*^(0)^ to the state (*n* – 1)^(0)^ is given by

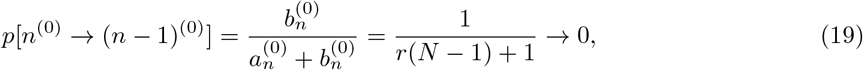

for *N* ≫ 1. Then it seems reasonable to neglect such transitions and to consider an effective discrete-state stochastic scheme as shown in Fig. 2c. Thus, we assume that 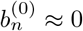 for *n* ≥ 2, and only the backward transition from the the state 1^(0)^ is assumed to be non-zero 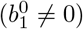.

The fixation dynamics in the approximate model can be explicitly analyzed as shown in the Supporting Information. More specifically, for fixation probabilities it is found that

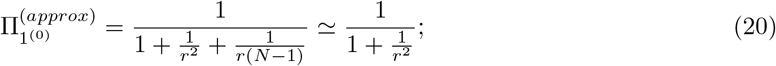

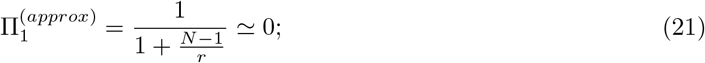

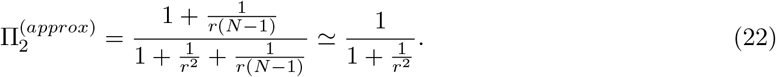

In addition, 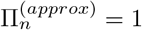 for *n* ≥ 3 and 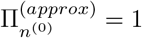 for *n* ≥ 2. Fig. 5 compares the predictions for the fixation probabilities for the approximate model of evolutionary dynamics. One can see that our approximation works quite well when *r*^2^ ≫ 1, and increasing the size of the system only slightly improves the agreement (Fig. 5a). But in all situations, the difference is only few percents between exact and approximate estimates of the fixation probabilities. At the same time, increasing the fitness parameter of the mutated cells has a much stronger effect (Fig. 5b). This is because the relation on which our approximation is based, Eq. (19), works even better for larger fitness parameters *r*. In all cases, we overestimate the fixation probabilities in comparison with exact expressions. This can be easily understood by again exploring the stochastic schemes in Figs. 2b and 2c. One can clearly see that our approximate model neglects the occasional backward steps in the upper chain of states that should only lower the fixation probability, in agreement with our predictions.

**Figure 5.**
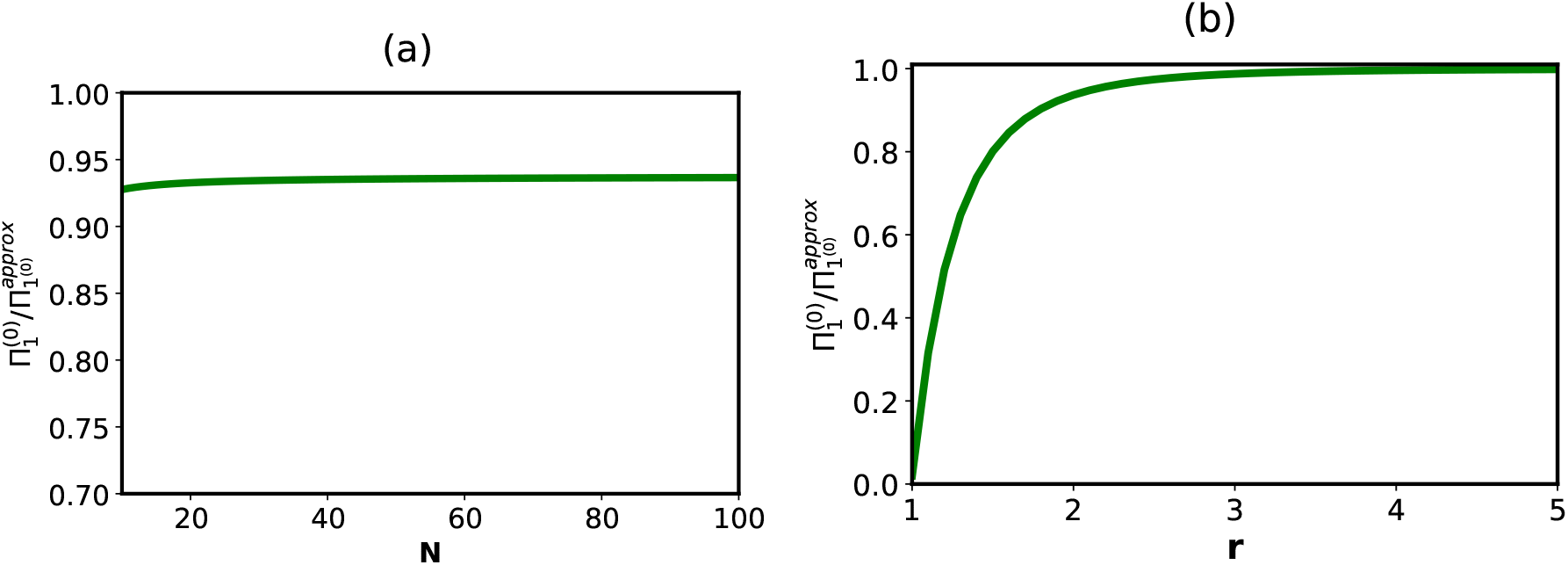
Comparison of fixation probabilities for the approximate and full models of evolutionary dynamics on star networks. Ratio of fixation probabilities a) as a function of the system size for *r* = 2, and b) as a function of the fitness parameter *r* for *N* = 100.

As shown in the Supporting Information, we can obtain the explicit expressions for the fixation times of the approximate model. It is found that

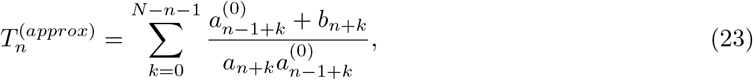

for *n* ≥ 3. In the limit *N* → ∞, it can be shown that 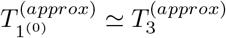 that eventually leads to the following estimate of the fixation time,

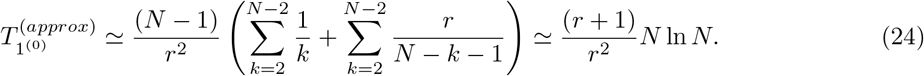

Thus, the approximate model correctly reproduces the scaling dependence of the fixation time [29].

Theoretical predictions for the fixation dynamics in the approximate and full models of evolutionary dynamics on star networks are presented in Fig. 6. One can see from Fig. 6a that approximating the fixation times is reasonable, although not as good as approximating the fixation probabilities: deviation of ~ 20% for times in comparison with ~ 5% for the probabilities for *r* = 2 case. The increasing the size of the system also does not have much effect. At the same time, increasing the advantage of the mutated cells (larger values of *r*) significantly improves the approximation: see Fig. 6b. As before, these observations can be understood by utilizing the discrete-state stochastic schemes from Fig. 2. Because the approximate model neglects the backward transitions from the states *n*^(0)^, it underestimates the fixation times by neglecting the backward and loop trajectories that should significantly slow down the overall dynamics in the system.

**Figure 6.**
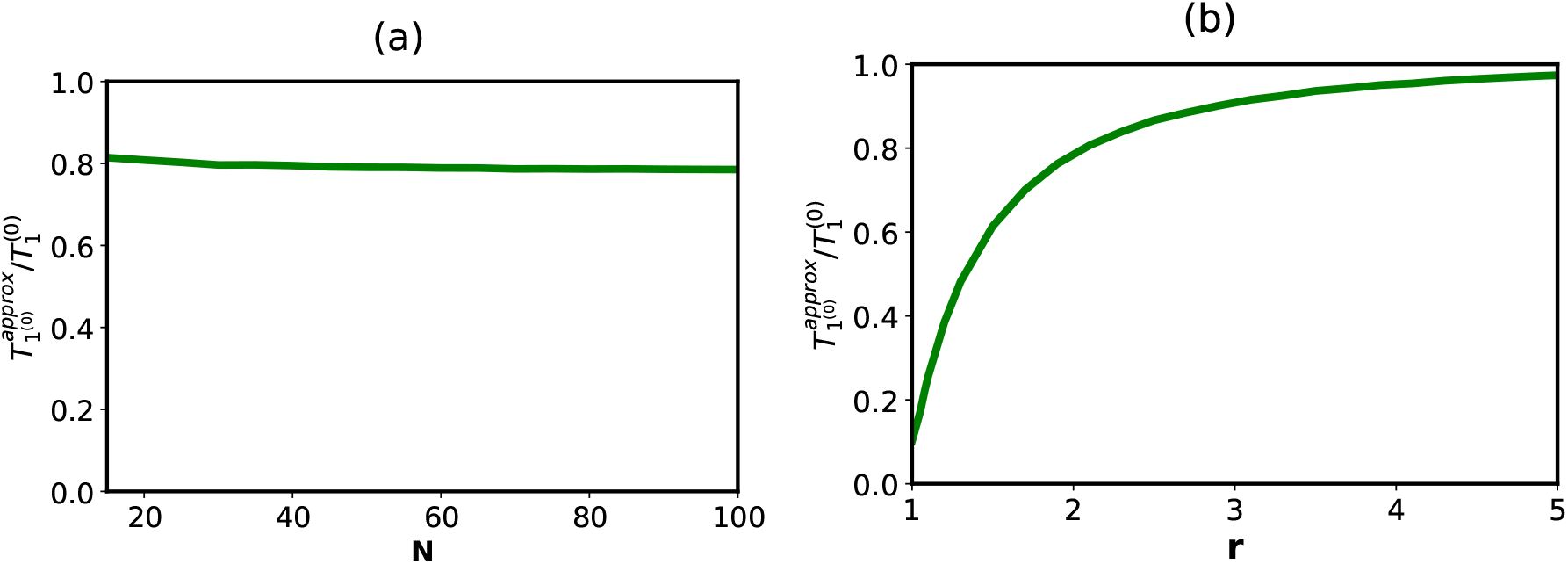
Comparison of fixation times for the approximate and full models of evolutionary dynamics on star networks. Ratio of fixation times a) as a function of the system size for *r* = 2, and b) as a function of the fitness parameter *r* for *N* =100.

Although the approximate model does not perfectly describe the fixation dynamics on the star network, it is valuable because it emphasizes better the main features of the evolutionary processes in these complex systems. The fixation amplification occurs only because the topological features of the system prevent the mutation elimination, while the fixation dynamics is quite slow because the system is frequently trapped in the unproductive states that are not on the pathway to the fixation. It is important to point out that this clear microscopic picture emerges as the result of mapping the evolutionary dynamics into the set of stochastic transitions between states specified by different numbers of mutated cells.

Another advantage of our discrete-state stochastic approach is that looking at the evolutionary processes as a motion in the effective free-energy landscape allows us to describe better the microscopic mechanisms of underlying processes and to discuss possible ways to optimize the evolution. This is schematically shown in Fig. 7. The fixation probability might be associated with the “free-energy” difference between the final state (fixation) and the initial state (one mutated cell), while the fixation times are given by the highest barrier on the pathway from the initial to the final states. For the well-mixed homogeneous system the advantage of reaching the fixation state is relatively modest, but the dynamics is also relatively fast. The situation is completely different for the evolutionary dynamics on the star network: see Fig. 7. Here, the advantage of reaching the fixation state is significant, but the “free-energy” barrier to accomplish this task is also quite large. As was discussed above, it is the consequence of topological properties of the star network. Thus, to accelerate the evolution, changes must be made to decrease these barriers and not in trying to increase the amplification of fixation probabilities. Our theoretical method suggest that one can concentrate on specific discrete-states where such changes will be the most effective.

**Figure 7.**
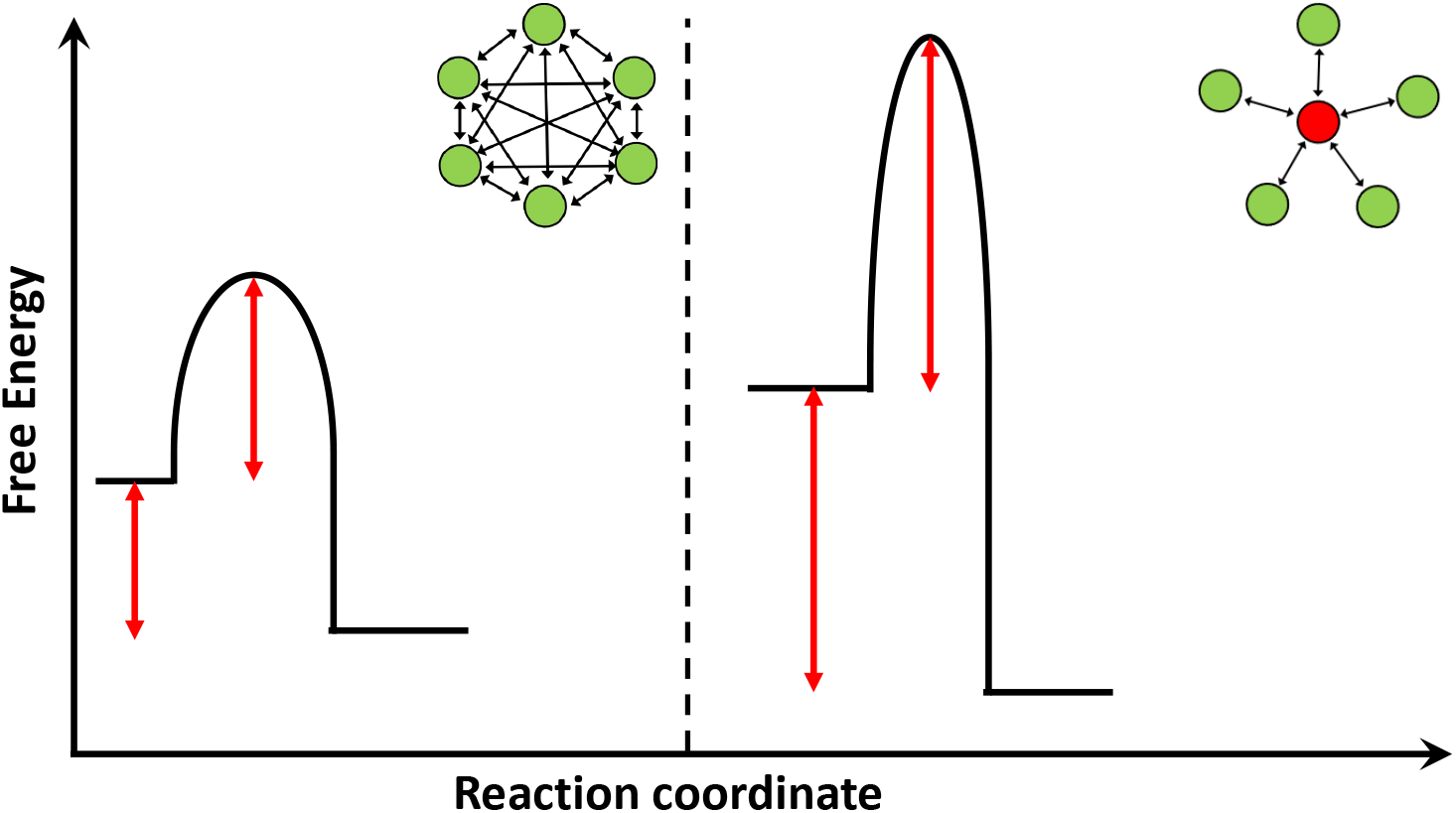
The effective free energy landscape for comparing evolutionary dynamics of full Moran model and star network.

## 3 Summary and Conclusions

In this paper, we developed a theoretical framework to investigate the role of inhomogeneity in evolutionary dynamics of structured populations. By analyzing the mutation fixation processes on the star network, we constructed a discrete-state stochastic model that provides a comprehensive description of the dynamics. Using the method of first-passage probabilities together with reasonable approximations explicit expressions for the fixation probabilities and fixation times are obtained. The presented theoretical method allowed us to better understand the microscopic origin of fixation amplification that is accompanied by significant increase in fixation times. It is argued that the amplification is the result of decreasing the probability of mutation elimination, but it does not increase the number of pathways to reach the fixation state, leading to slowing down in the fixation dynamics.

The mapping of evolutionary dynamics on inhomogeneous networks into the motion in the effective free-energy landscape provides new insights on the mechanisms of these complex processes. It also suggests how these systems can be modified to optimize the evolutionary output.

The important advantage of our theoretical approach is that the method can be extended for studying the evolutionary processes on other inhomogeneous systems. Specifically, we plan to generalize our theoretical arguments for analyzing the evolutionary dynamics on *l*-star networks where there are *l* star nodes that are connected with all *N* – l branched cells. The system considered in this paper is a special case with *l* = 1. It will be important to understand how the degree of amplification in those systems correlates with the fixation times. In addition, our theoretical method can be extended for dynamic networks where topological features might fluctuate between several different arrangements. It will be also important to apply these theoretical results for understanding cancer initiation and tumor formation.

## Data accessibility

This article has no additional data.

## Author contributions

H.T. and A.B.K designed the research; H.T., C.L., D.S.K. and C.S performed the research; H.T., C.L. and A.B.K wrote the manuscript.

## Acknowledgements

We acknowledge the support from the Welch Foundation (C-1559), from the NSF (CHE-1953453 and MCB-1941106), and from the Center for Theoretical Biological Physics sponsored by the NSF (PHY-2019745).

## Competing Interests

We declare that we have no competing interests.

## Funding Statement

The work was supported by the Welch Foundation (C-1559), by the NSF (CHE-1953453 and MCB-1941106), and by the Center for Theoretical Biological Physics sponsored by the NSF (PHY-2019745).

## Notes

### Competing Interest Statement

The authors have declared no competing interest.

