## Supporting information for "Theoretical understanding of evolutionary dynamics on inhomogeneous networks"

February 2, 2023

In this Supporting Information, we present the derivations of equations that are discussed in the main text of the paper.

#### 1 Calculations of Transition Rates

To calculate transition rates, we consider each birth/death Moran process as a two-step process, as shown in Fig. S1 [see also Ref. [1]]. Here we assume that the additional cells are removed with a rate $A \gg r$. The transition rates are then given by,

$$a_n = \frac{nr\frac{A(N-n)}{N-1}}{nr + A(n-1) + \frac{A(n-1)}{N-1} + \frac{A(N-n)}{N-1}} \simeq \frac{r(N-n)}{N-1} \tag{1}$$

$$b_n^{(0)} = \frac{\frac{An(N-n)}{N-1}}{N-n+\frac{An}{N-1} + A(N-n-1) + \frac{A(N-n-1)}{N-1}} \simeq \frac{n}{N-1} \tag{2}$$

$$a_n^{(0)} = \frac{Arn}{rn+A} \simeq nr \tag{3}$$

$$b_n = \frac{A(N-n)}{A+nr+N-n} \simeq N-n \tag{4}$$

#### Exact Solution for the Star Network Model with $N = 3$ Cells

In this section, we derive a full solution for the specific model of evolutionary dynamics with just $N = 3$ cells. As shown in the Fig. ??, there are 4 possible states in this system. The

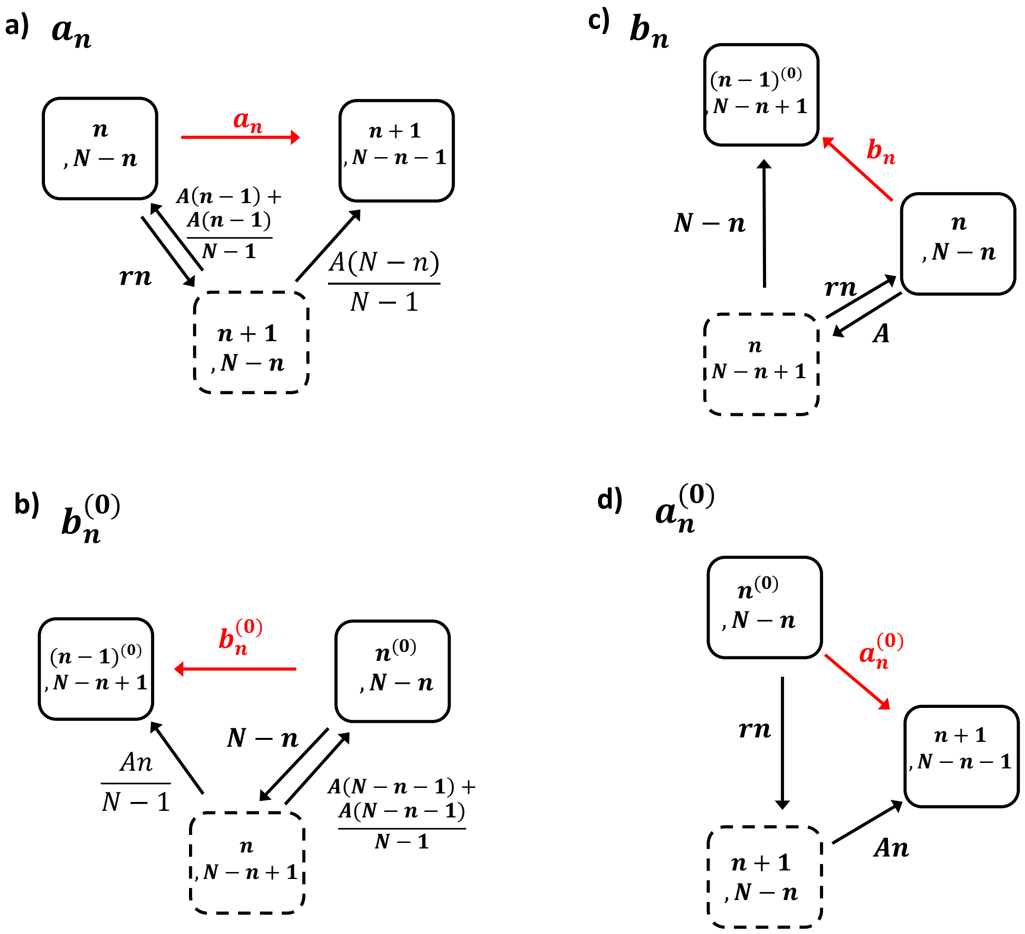


Figure S1: Schematic view for the calculation of transition rates: a) $a_n$ b) $b_n^{(0)}$ c) $b_n$, and d) $a_n^{(0)}$. Note that for transition shown in a) and d), the intermediate states contain $n+1$ mutated cells and $N-n$ normal cells, while for transitions in b) and c), the intermediate states contain $n$ mutated cells and $N-n+1$ normal cells.

temporal evolution of the first-passage probabilities is given by,

$$\frac{dF_1^{(0)}}{dt} = a_1^{(0)} F_2 - (a_1^{(0)} + b_1^{(0)}) F_1^{(0)} \tag{5}$$

$$\frac{dF_2^{(0)}}{dt} = a_2^{(0)} F_{fix} + b_2^{(0)} F_1^{(0)} - (a_2^{(0)} + b_2^{(0)}) F_2^{(0)} \tag{6}$$

$$\frac{dF_1}{dt} = a_1 F_2 - (a_1 + b_1) F_1 \tag{7}$$

$$\frac{dF_2}{dt} = a_2 F_{fix} + b_2 F_1^{(0)} - (a_2 + b_2) F_2, \tag{8}$$

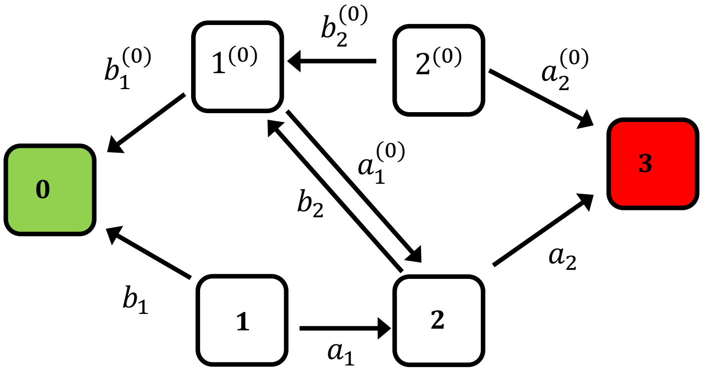


Figure S2: Schematic view of the discrete-state stochastic model of star-structured population for $N = 3$.

where $F_{fix}(t) = \delta(t)$. Performing Laplace transformation, we get

$$(s + a_1^{(0)} + b_1^{(0)})\widetilde{F_1^{(0)}} = a_1^{(0)}\widetilde{F_2} \tag{9}$$

$$(s + a_2^{(0)} + b_2^{(0)})\widetilde{F_2^{(0)}} = a_2^{(0)}\widetilde{F_3} + b_2^{(0)}\widetilde{F_1^{(0)}} \tag{10}$$

$$(s + a_1 + b_1)\widetilde{F_1} = a_1\widetilde{F_2} \tag{11}$$

$$(s + a_2 + b_2)\widetilde{F_2} = a_2\widetilde{F_3} + b_2\widetilde{F_1^{(0)}} \tag{12}$$

This system of equations can be solved, yielding,

$$\widetilde{F_1^{(0)}} = -\frac{a_1a_2}{-(a_1^{(0)}a_2 - a_2b_1^{(0)} + a_1b_2 - a_1^{(0)}b_2 - b_1^{(0)}b_2 - a_1^{(0)}s - a_2s - b_1^{(0)}s - b_2s - s^2)} \tag{13}$$

$$\widetilde{F_1} = -\frac{a_2(a_1a_1^{(0)} + a_1b_1^{(0)} + a_1s)}{-(a_1 + b_1 + s)(a_1^{(0)}a_2 - a_2b_1^{(0)} + a_1b_2 - a_1^{(0)}b_2 - b_1^{(0)}b_2 - a_1^{(0)}s - a_2s - b_1^{(0)}s - b_2s - s^2)} \tag{14}$$

From these expressions, one can calculate the fixation probabilities,

$$\Pi_1 = \widetilde{F_1}(s = 0) = -\frac{a_2(a_1a_1^{(0)} + a_1b_1^{(0)})}{(a_1 + b_1)(-a_1^{(0)}a_2 - a_2b_1^{(0)} + a_1b_2 - a_1^{(0)}b_2 - b_1^{(0)}b_2)} \tag{15}$$

$$\Pi_1^{(0)} = \widetilde{F_1^{(0)}}(s = 0) = -\frac{a_1a_2}{-a_1^{(0)}a_2 - a_2b_1^{(0)} + a_1b_2 - a_1^{(0)}b_2 - b_1^{(0)}b_2} \tag{16}$$

In Fig. 3, we compare these results with Monte-Carlo computer simulations, and excellent agreement is found for all sets of parameters.

Also, fixation times can be evaluated as

$$T_1^{(0)} = -\frac{\frac{\partial \widetilde{F_1^{(0)}}}{\partial s}|_{s=0}}{\Pi_1^{(0)}} = -\frac{1}{\Pi_1^{(0)}}\left[\frac{a_1a_2(a_1^{(0)} + a_2 + b_1^{(0)} + b_2)}{(a_1^{(0)}a_2 + a_2b_1^{(0)} - a_1b_2 + a_1^{(0)}b_2 + b_1^{(0)}b_2)^2}\right]; \tag{17}$$

and

$$T_1 = -\frac{\frac{\partial \widetilde{F_1}}{\partial s}|_{s=0}}{\Pi_1} = \frac{1}{\Pi_1}\left[-\frac{a_2(a_1a_1^{(0)}+a_1b_1^{(0)})(a_1^{(0)}+a_2+b_1^{(0)}+b_2)}{(a_1+b_1)(-a_1^{(0)}a_2-a_2b_1^{(0)}+a_1b_2-a_1^{(0)}b_2-b_1^{(0)}b_2)^2}\right]+ \frac{1}{\Pi_1}\left[\frac{a_1a_2}{(a_1+b_1)(-a_1^{(0)}a_2-a_2b_1^{(0)}+a_1b_2-a_1^{(0)}b_2-b_1^{(0)}b_2)}\right]+ \frac{1}{\Pi_1}\left[\frac{a_2(a_1a_1^{(0)}+a_1b_1^{(0)})}{(a_1+b_1)^2(-a_1^{(0)}a_2-a_2b_1^{(0)}+a_1b_2-a_1^{(0)}b_2-b_1^{(0)}b_2}\right]. \quad (18)$$

Fig. 4 compares our predictions for the fixation times with computer simulations, and again excellent agreement is found for all ranges of parameters.

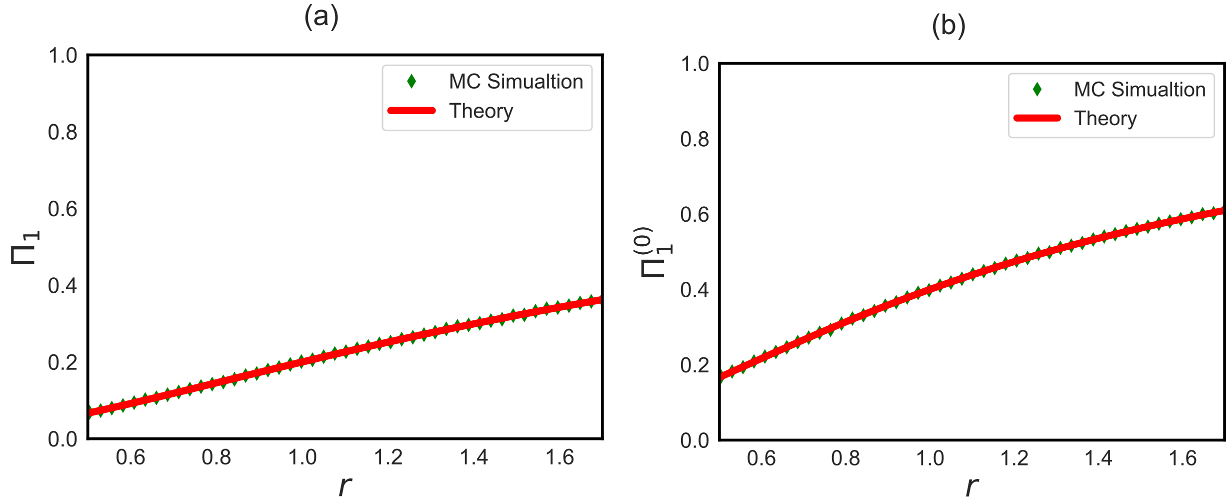


Figure S3: Comparison of approximate result with Monte Carlo computer simulation for a) $\Pi_1^{(0)}$ and b) $\Pi_2$

### 2 Calculations of Fixation Probabilities for Arbitrary Size Star Networks

As described in the main text, we define $F_n(t)$ ($F_n^{(0)}(t)$) as the probability density of reaching state $N$ at time $t$ if at $t = 0$ the system started in the state $n$ ($n^{(0)}$). A schematic picture for the model is shown in Fig. 2(b) in the main text. Temporal evolution of these first-passage

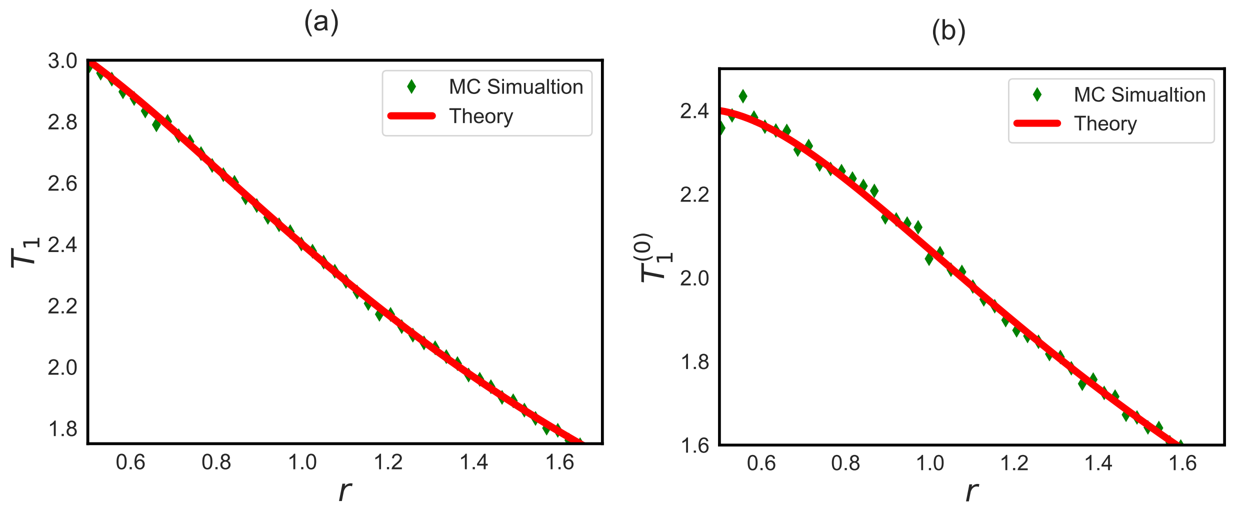


Figure S4: Comparison of approximate result with Monte Carlo computer simulation for a) $T_1$ and b) $T_1^{(0)}$.

probabilities can be described as a set of backwards master equations,

$$\frac{dF_1^{(0)}}{dt} = a_1^{(0)} F_2 - (a_1^{(0)} + b_1^{(0)}) F_1^{(0)}, \tag{19}$$

$$\frac{dF_1}{dt} = a_1 F_2 - (a_1 + b_1) F_1, \tag{20}$$

$$\frac{dF_{N-1}^{(0)}}{dt} = b_{N-1}^{(0)} F_{N-2}^{(0)} + a_{N-1}^{(0)} F_{fix} - (a_{N-1}^{(0)} + b_{N-1}^{(0)}) F_{N-1}^{(0)}, \tag{21}$$

$$\frac{dF_{N-1}}{dt} = b_{N-1} F_{N-2}^{(0)} + a_{N-1} F_{fix} - (a_{N-1} + b_{N-1}) F_{N-1}, \tag{22}$$

$$\frac{dF_n^{(0)}}{dt} = a_n^{(0)} F_{n+1} + b_n^{(0)} F_{n-1}^{(0)} - (a_n^{(0)} + b_n^{(0)}) F_n^{(0)}, \tag{23}$$

$$\frac{dF_n}{dt} = a_n F_{n+1} + b_n F_{n-1}^{(0)} - (a_n + b_n) F_n, \tag{24}$$

where $F_{fix}(t)$ corresponds to the state in which all cells are mutated. The initial condition is again $F_{fix} = \delta(t)$. To proceed further, we perform Laplace transformation over backward master equations,

$$(s + a_1^{(0)} + b_1^{(0)}) \widetilde{F_1^{(0)}} = a_1^{(0)} \widetilde{F_2} \tag{25}$$

$$(s + a_1 + b_1) \widetilde{F_1} = a_1 \widetilde{F_2} \tag{26}$$

$$(s + a_{N-1}^{(0)} + b_{N-1}^{(0)}) \widetilde{F_{N-1}^{(0)}} = b_{N-1}^{(0)} \widetilde{F_{N-2}^{(0)}} + a_{N-1}^{(0)} \tag{27}$$

$$(s + a_{N-1} + b_{N-1}) \widetilde{F_{N-1}} = b_{N-1} \widetilde{F_{N-2}^{(0)}} + a_{N-1} \tag{28}$$

$$(s + a_n^{(0)} + b_n^{(0)}) \widetilde{F_n^{(0)}} = a_n^{(0)} \widetilde{F_{n+1}} + b_n^{(0)} \widetilde{F_{n-1}^{(0)}} \tag{29}$$

$$(s + a_n + b_n) \widetilde{F_n} = a_n \widetilde{F_{n+1}} + b_n \widetilde{F_{n-1}^{(0)}} \tag{30}$$

We assume that at the steady state (which corresponds to $s \to 0$) $\widetilde{F_1^{(0)}}(s)$ and $\widetilde{F_1}(s)$ can be expanded as follows,

$$\widetilde{F_1^{(0)}} \simeq \Pi_n^{(0)} - \Pi_n^{(0)} T_n^{(0)} s + ... \tag{31}$$

$$\widetilde{F_1} \simeq \Pi_n - \Pi_n T_n s + ... \tag{32}$$

Substituting Eqs. 31 and 32 into Laplace transformation of backward master equations, we get the following relations for the fixation probabilities,

$$(a_1^{(0)} + b_1^{(0)})\Pi_1^{(0)} = a_1^{(0)}\Pi_2 \tag{33}$$

$$(a_1 + b_1)\Pi_1 = a_1\Pi_2 \tag{34}$$

$$(a_{N-1}^{(0)} + b_{N-1}^{(0)})\Pi_{N-1}^{(0)} = b_{N-1}^{(0)}\Pi_{N-2}^{(0)} + a_{N-1}^{(0)} \tag{35}$$

$$(a_{N-1} + b_{N-1})\Pi_{N-1} = b_{N-1}\Pi_{N-2}^{(0)} + a_{N-1} \tag{36}$$

$$(a_n^{(0)} + b_n^{(0)})\Pi_n^{(0)} = a_n^{(0)}\Pi_{n+1} + b_n^{(0)}\Pi_{n-1}^{(0)} \tag{37}$$

$$(a_n + b_n)\Pi_n = a_n\Pi_{n+1} + b_n\Pi_{n-1}^{(0)} \tag{38}$$

These equations can be solved using boundary conditions $\Pi_N = 1$ and $\Pi_0^{(0)} = 0$. Equations (37) and (38) can be rewritten:

$$\lambda\Pi_n^{(0)} = \Pi_{n+1} + (\lambda - 1)\Pi_{n-1}^{(0)} \tag{39}$$

$$\gamma\Pi_n = \Pi_{n+1} + (\gamma - 1)\Pi_{n-1}^{(0)} \tag{40}$$

where $\gamma$ and $\lambda$ are defined below:

$$\gamma = 1 + \frac{b_n}{a_n} = 1 + \frac{N-1}{r} \tag{41}$$

$$\lambda = 1 + \frac{b_n^{(0)}}{a_n^{(0)}} = 1 + \frac{1}{r(N-1)} \tag{42}$$

We are looking for solutions in the following form,

$$\Pi_n^{(0)} = C + Ax^n, \tag{43}$$

$$\Pi_{n+1} = C + Bx^n. \tag{44}$$

Substituting into the Eqs. 39 and 40 yields,

$$\lambda Ax^n = Bx^n + (\lambda - 1)Ax^{n-1}, \tag{45}$$

$$\gamma Bx^{n-1} = Bx^n + (\gamma - 1)Ax^{n-1}. \tag{46}$$

Now, solving the equations for $x$ produces

$$x = \frac{(\lambda - 1)(\frac{A}{B})}{\lambda\frac{A}{B} - 1}, \tag{47}$$

$$x = \gamma - (\gamma - 1)(\frac{A}{B}). \tag{48}$$

To proceed further, we define a new function $y = \frac{A}{B}$. Thus, Eq. 47 leads to the following quadratic equation for $y$,

$$\lambda(\gamma-1)y^2 - y[\lambda(\gamma-1)+\gamma] + \gamma = 0. \tag{49}$$

Solving for $y$ gives us $y_1 = 1$ and $y_2 = \frac{\gamma}{\lambda(\gamma-1)}$. For $y = 1$ we get $x = 1$, which is not acceptable. Thus, physically acceptable root is only $y = \frac{\gamma}{\lambda(\gamma-1)}$. Thus, $x$ takes the form,

$$x = \frac{\gamma}{\lambda}(\lambda - 1). \tag{50}$$

.

Using boundary conditions, $\Pi_0 = 0$, we obtain,

$$C = -A, \tag{51}$$

$$B = \frac{A}{y}. \tag{52}$$

Furthermore, using $\Pi_N = 1$, we derive,

$$1 = -A + Bx^{N-1} = -A + \frac{A}{y}x^{N-1}. \tag{53}$$

After some algebra, it leads to

$$A = \frac{1}{\frac{\lambda}{\gamma}(\gamma-1)x^{N-1} - 1}. \tag{54}$$

Now, having all constants $A, B, C, x$ to be explicitly expressed allows us to calculate $\Pi_1$ and $\Pi_1^{(0)}$,

$$\Pi_1^{(0)} = A(x-1) = \frac{1 - \frac{1}{r^2}}{\left[1 + \frac{1}{r(N-1)}\right]\left[1 - \frac{1}{r^2}\left(\frac{1+\frac{N-1}{r}}{1+r(N-1)}\right)^{N-2}\right]}; \tag{55}$$

$$\Pi_1 = C + B = \frac{1 - \frac{1}{r^2}}{\left[1 + \frac{N-1}{r}\right]\left[1 - \frac{1}{r^2}\left(\frac{1+\frac{N-1}{r}}{1+r(N-1)}\right)^{N-2}\right]}; \tag{56}$$

and for general $n$ we have

$$\Pi_n^{(0)} = \frac{1 + \frac{1}{r(N-1)} - \left(\frac{1}{r^2} + \frac{1}{r(N-1)}\right)^n}{\left[1 + \frac{1}{r(N-1)}\right]\left[1 - \frac{1}{r^2}\left(\frac{1+\frac{N-1}{r}}{1+r(N-1)}\right)^{N-2}\right]}. \tag{57}$$

$$\Pi_n = \frac{1 - \frac{1}{r^2}\left(\frac{1+\frac{N-1}{r}}{1+r(N-1)}\right)^{n-2}}{\left[1 - \frac{1}{r^2}\left(\frac{1+\frac{N-1}{r}}{1+r(N-1)}\right)^{N-2}\right]}. \tag{58}$$

### 3 Approximate Model

The schematic view for the approximate model is shown in Fig. 2c in the main text. The dynamical evolution of the system is governed by the following forward master equations,

$$\frac{dF_1^{(0)}}{dt} = a_1^{(0)} F_2 - (a_1^{(0)} + b_1^{(0)}) F_1^{(0)}, \tag{59}$$

$$\frac{dF_{N-1}}{dt} = b_{N-1} F_{N-2}^{(0)} + a_{N-1} F_{fix} - (a_{N-1} + b_{N-1}) F_{N-1}, \tag{60}$$

$$\frac{dF_n^{(0)}}{dt} = a_n^{(0)} F_{n+1} - a_n^{(0)} F_n^{(0)}, \tag{61}$$

$$\frac{dF_n}{dt} = a_n F_{n+1} + b_n F_{n-1}^{(0)} - (a_n + b_n) F_n, \tag{62}$$

with initial condition $F_{fix}(t) = \delta(t)$. Performing Laplace transformation yields,

$$(s + a_1^{(0)} + b_1^{(0)}) \widetilde{F_1^{(0)}} = a_1^{(0)} \widetilde{F_2} \tag{63}$$

$$(s + a_{N-1} + b_{N-1}) \widetilde{F_{N-1}} = b_{N-1} \widetilde{F_{N-2}^{(0)}} + a_{N-1} \tag{64}$$

$$(s + a_n^{(0)}) \widetilde{F_n^{(0)}} = a_n^{(0)} \widetilde{F_{n+1}} \tag{65}$$

$$(s + a_n + b_n) \widetilde{F_n} = a_n \widetilde{F_{n+1}} + b_n \widetilde{F_{n-1}^{(0)}} \tag{66}$$

Again, we use the Taylor expansions:

$$\widetilde{F_n^{(0)}}(s) \approx \Pi_n^{(0)} - \Pi_n^{(0)} T_n^{(0)} s \tag{67}$$

$$\widetilde{F_n(s)} \approx \Pi_n - \Pi_n T_n s \tag{68}$$

Let us start with evaluating fixation probabilities. Using Fig. 2c in the main text, it is easy to show that

$$\Pi_n = 1 \tag{69}$$

for $n = 3, 4, ..., N-1$, and

$$\Pi_n^{(0)} = 1 \tag{70}$$

for $n = 2, 3, 4, ..., N-2$. However, for $n = 1, 2$ substituting Eqs. 67 into Eqs. 67 produces

$$(a_2 + b_2)\Pi_2 = a_2 \Pi_3 + b_2 \Pi_1^{(0)} \tag{71}$$

$$(a_1^{(0)} + b_1^{(0)}) \Pi_1^{(0)} = a_1^{(0)} \Pi_2 \tag{72}$$

Solving these equations gives us,

$$\Pi_1^{(0)} = \frac{a_2 a_1^{(0)}}{a_2 a_1^{(0)} + a_2 b_1^{(0)} + b_2 b_1^{(0)}}, \tag{73}$$

$$\Pi_2 = \frac{a_2 a_1^{(0)} + a_2 b_1^{(0)}}{a_2 a_1^{(0)} + a_2 b_1^{(0)} + b_2 b_1^{(0)}}. \tag{74}$$

$$\tag{75}$$

Using the explicit expressions for the transitions rates, we obtain

$$\Pi_1^{(0)} = \frac{1}{1+\frac{1}{r^2}+\frac{1}{r(N-1)}} \simeq \frac{1}{1+\frac{1}{r^2}} \tag{76}$$

$$\Pi_2 = \frac{1+\frac{1}{r(N-1)}}{1+\frac{1}{r^2}+\frac{1}{r(N-1)}} \simeq \frac{1}{1+\frac{1}{r^2}}. \tag{77}$$

Now, we can calculate fixation times by substituting Eqs. 67 into Eqs. 67. After simplification, we get the following set of equations,

$$\Pi_1^{(0)} - \Pi_1^{(0)} T_1^{(0)} (a_1^{(0)} + b_1^{(0)}) = -a_1^{(0)} \Pi_2 T_2, \tag{78}$$

$$\Pi_{N-1} - \Pi_{N-1} T_{N-1} (a_{N-1} + b_{N-1}) = -b_{N-1} \Pi_{N-2}^{(0)} T_{N-2}^{(0)}, \tag{79}$$

$$\Pi_n^{(0)} - a_n^{(0)} T_n^{(0)} \Pi_n^{(0)} = -a_n^{(0)} \Pi_{n+1} T_{n+1}, \tag{80}$$

$$\Pi_n - \Pi_n T_n (a_n + b_n) = -a_n \Pi_{n+1} T_{n+1} - b_n \Pi_{n-1}^{(0)} T_{n-1}^{(0)}. \tag{81}$$

For $n \geq 3$, these equations can be further simplified into

$$1 - T_1^{(0)} (a_1^{(0)} + b_1^{(0)}) = -a_1^{(0)} \frac{\Pi_2}{\Pi_1^{(0)}} T_2 \tag{82}$$

$$1 - T_{N-1} (a_{N-1} + b_{N-1}) = -b_{N-1} T_{N-2}^{(0)} \tag{83}$$

$$1 - T_n^{(0)} a_n^{(0)} = -a_n T_{n+1} \tag{84}$$

$$1 - T_n (a_n + b_n) = -a_n T_{n+1} - b_n T_{n-1}^{(0)} \tag{85}$$

$$\Pi_2 - \Pi_2 T_2 (a_2 + b_2) = -a_2 T_3 - b_2 \Pi_1^{(0)} T_1^{(0)} \tag{86}$$

After some algebra we derive

$$T_{N-2}^{(0)} = \frac{a_{N-2}^{(0)} + a_{N-1} + b_{N-1}}{a_{N-1} a_{N-2}^{(0)}}, \tag{87}$$

$$T_{N-1} = \frac{a_{N-2}^{(0)} + b_{N-1}}{a_{N-1} a_{N-2}^{(0)}}, \tag{88}$$

$$T_{n-1}^{(0)} = T_{n+1} + \frac{a_{n-1}^{(0)} + a_n + b_n}{a_n a_{n-1}^{(0)}}, \tag{89}$$

$$T_n = T_{n+1} + \frac{a_{n-1}^{(0)} + b_n}{a_n a_{n-1}^{(0)}}. \tag{90}$$

For $n \geq 3$, we get the following expression for the fixation times,

$$T_n = \sum_{k=0}^{N-n-1} \frac{a_{n-1+k}^{(0)} + b_{n+k}}{a_{n+k} a_{n-1+k}^{(0)}}. \tag{91}$$

However, we are interested in $T_1^{(0)}$. Using Eqs. 82 and 86 we obtain,

$$T_1^{(0)} = \left(1 + \frac{b_1^{(0)}}{a_1^{(0)}} - \frac{b_2}{a_2 + b_2}\right)^{-1} \frac{1}{\Pi_1^{(0)}} \left(\frac{\Pi_1^{(0)}}{a_1^{(0)}} + \frac{\Pi_2}{a_2 + b_2} + \frac{a_2}{a_2 + b_2} T_3\right). \tag{92}$$

For $N \to \infty$, Eq. 92 simplifies into

$$T_1^{(0)} \simeq T_3. \tag{93}$$

After some algebra, it can bhe shown that the fixation times are given by

$$T_1^{(0)} \simeq T_3 = \frac{N-1}{r^2} \sum_{k=2}^{N-2} \frac{1}{k} + \frac{N-1}{r^2} \sum_{k=2}^{N-2} \frac{r}{N-k-1}. \tag{94}$$

One can evaluate the summation by integration,

$$\sum_{k=a}^{b} \frac{1}{k} \simeq \int_a^b \frac{1}{x} dx \simeq \ln \frac{b}{a} \tag{95}$$

Thus for $N \to \infty$ , the fixation time is estimated as

$$T_1^{(0)} = \frac{r+1}{r^2} N \ln(N). \tag{96}$$
